## Supplementary Figures S1-9 for "Sorting of secretory proteins at the trans-Golgi network by human TGN46"

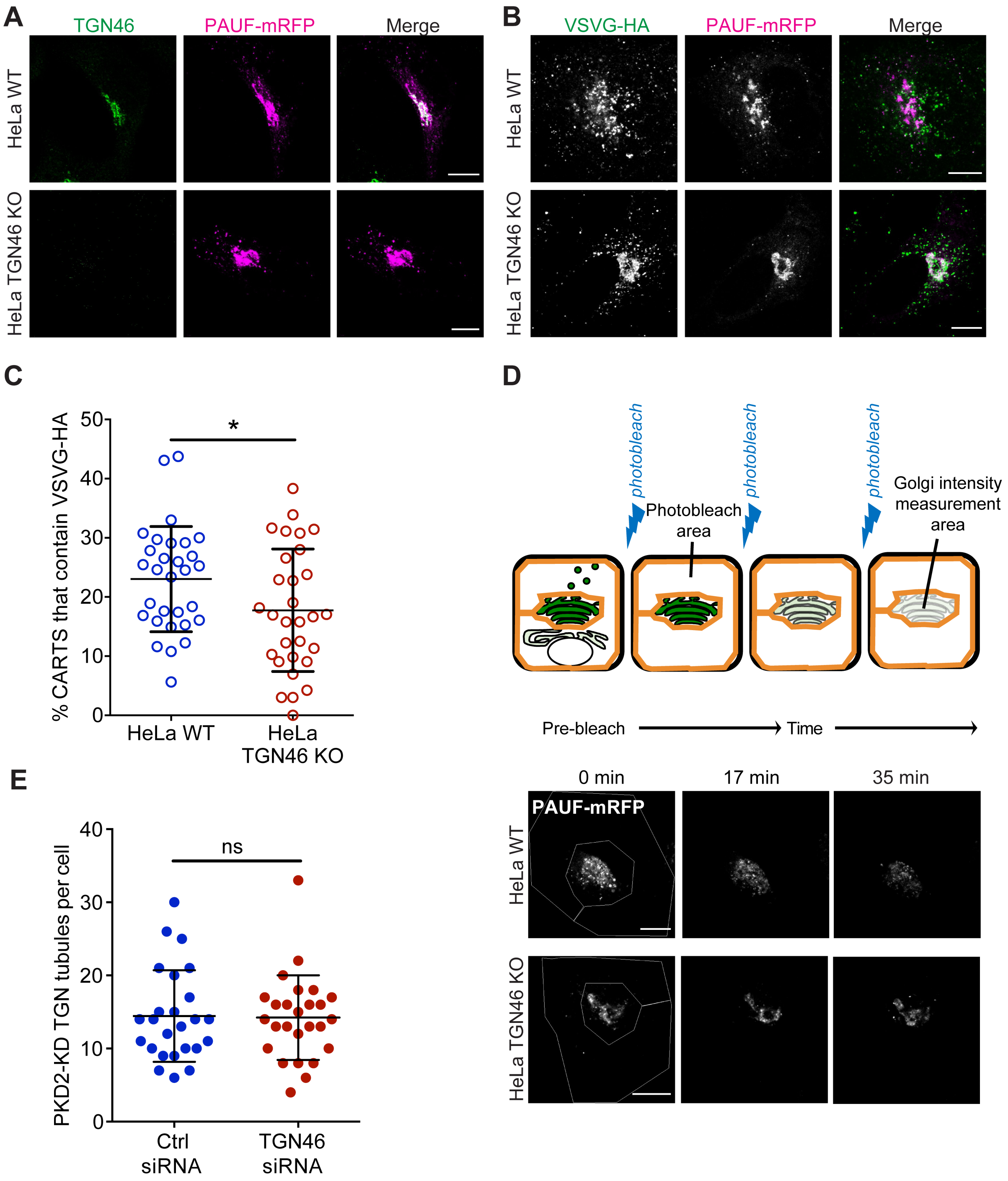


**Figure S1. TGN46 is required for cargo sorting and loading into CARTS. (A)** HeLa WT or HeLa TGN46-KO cells expressing PAUF-mRFP were fixed, processed for immunostaining, and the localization of endogenous TGN46 and PAUF-mRFP was monitored by immunofluorescence microscopy. **(B)** HeLa WT or HeLa TGN46-KO cells expressing VSVG-HA and PAUF-mRFP were fixed, processed for immunostaining, and the localization of VSVG-HA and PAUF-mRFP was monitored by immunofluorescence microscopy. **(C)** Percentage of PAUF-mRFP-containing carriers (CARTS) that are also positive for VSVG-HA in HeLa WT or TGN46-KO cells, quantified from confocal micrographs as those shown in **(B)**. Results are from at least 10 cells from each of n=3 independent experiments (individual values shown, with mean ± stdev). Unpaired two-tailed t test (*, $p \leq 0.05$). **(D)** *Top:* Schematic representation of FLIP experiments (see text and methods for details). *Bottom:* Fluorescence microscopy images obtained from a characteristic FLIP experiment assessing the export rate of PAUF-mRFP from the perinuclear area in either HeLa WT or HeLa TGN46-KO cells. Time from the beginning of the FLIP experiments is indicated, and the area enclosed by the white lines shown in the left images denotes the photobleached area. **(E)** Quantification of the number of PKD2-KD-containing tubules per cell in HeLa cells transfected with control (Ctrl) or TGN46-targeting siRNA. Results are from at least 10 cells from each of n=3 independent experiments (individual values shown, with mean ± stdev). Unpaired two-tailed t test (ns, $p > 0.05$). Scale bars in **(A, B, D)** are 10 µm.

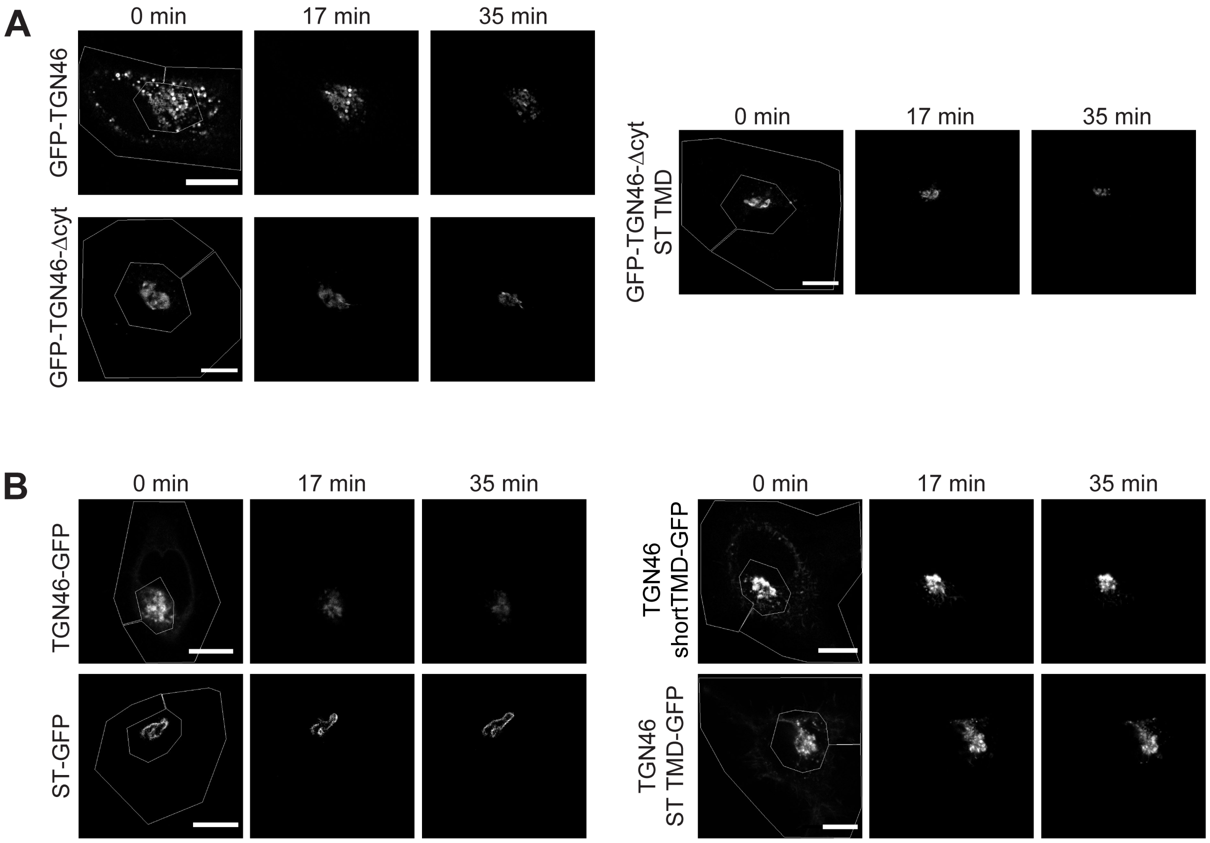


**Figure S2. FLIP experiments monitor Golgi residence times of different proteins. (C)** Fluorescence microscopy images obtained from a characteristic FLIP experiment assessing the export rate of GFP-TGN46 (top left row), GFP-TGN46-Δcyt (bottom left row), or GFP-TGN46-Δcyt-ST TMD (right row) from the perinuclear area in HeLa cells. Time from the beginning of the FLIP experiments is indicated, and the area enclosed by the white lines shown in the left images denotes the photobleached area. **(C)** Fluorescence microscopy images obtained from a characteristic FLIP experiment assessing the export rate of TGN46 WT-GFP (top left row), ST-GFP (bottom left row), TGN46-shortTMD-GFP (top right row), or TGN46-ST TMD-GFP (bottom right row) from the perinuclear area in HeLa cells. Time from the beginning of the FLIP experiments is indicated, and the area enclosed by the white lines shown in the left images denotes the photobleached area. Scale bars are 10 μm.

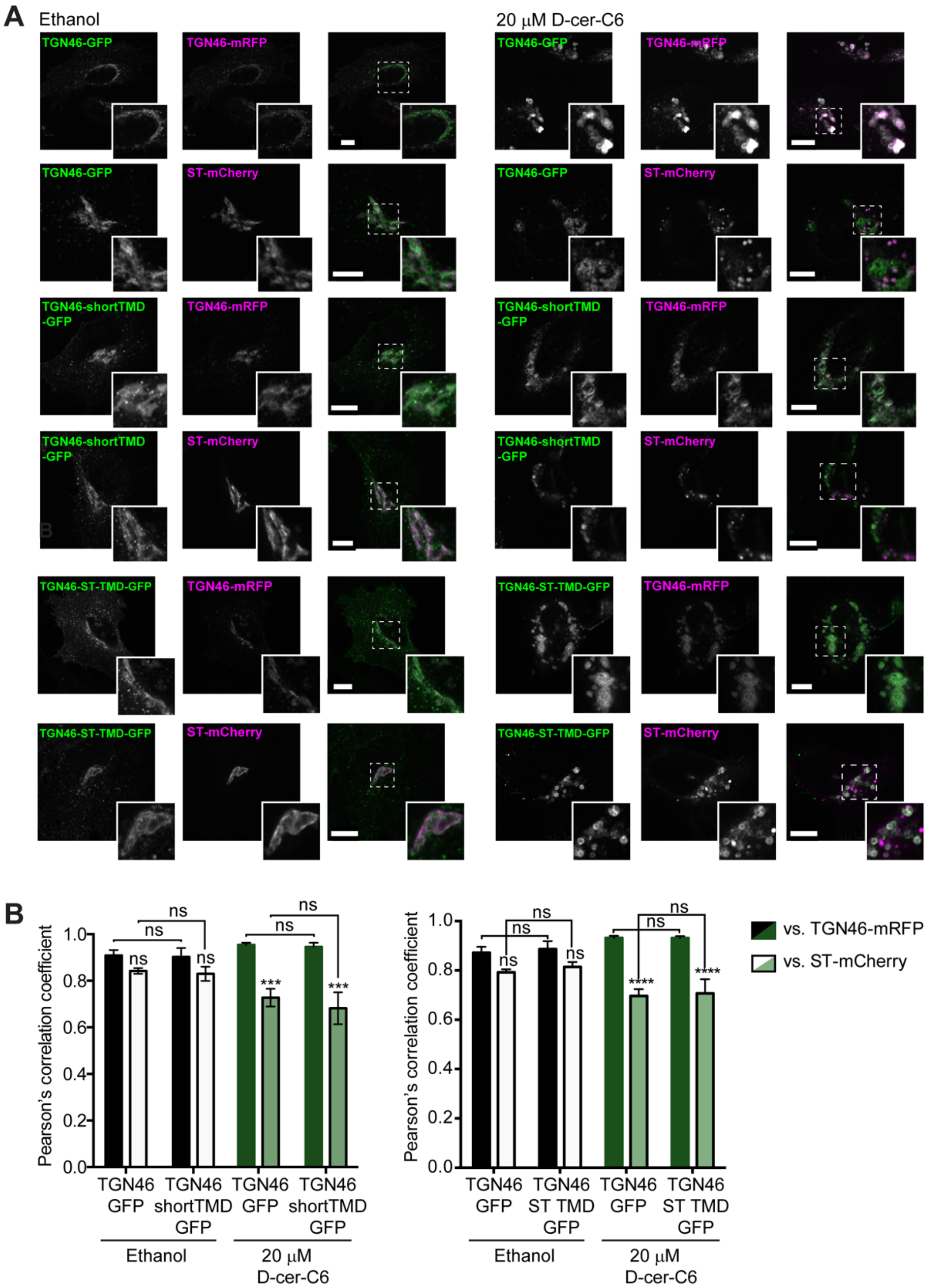


**Figure S3. Effect of sphingolipid metabolism on the intra-Golgi localization of TGN46 mutants. (A)** HeLa cells expressing TGN46-GFP, TGN46-shortTMD-GFP, or TGN46-ST TMD-GFP together with TGN46-mRFP or ST-mCherry were treated with ethanol or with 20 μM D-cer-C6 for 4 h. The localization of these proteins was monitored by fluorescence microscopy. Insets correspond to zoom-in areas of the dashed, white boxed areas. Scale bar, 10 μm. **(B)** Quantitation of the relative colocalization of the different proteins in the experiments shown in **(A)**, as measured by the Pearson's correlation coefficient between the green and red channels. Bars show the mean values ± s.e.m. of $\geq$10 cells counted from three independent experiments.

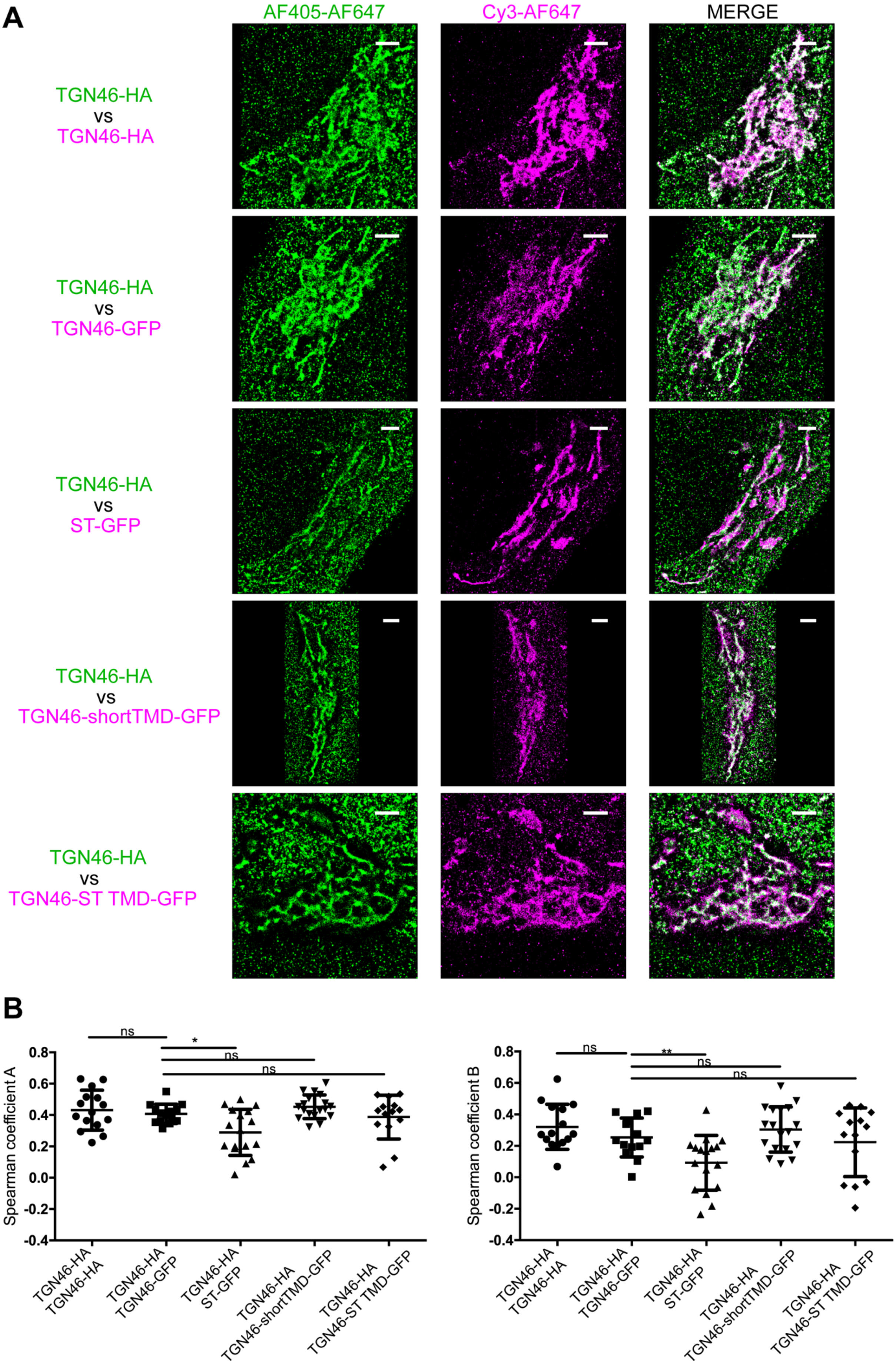

A
AF405-AF647
Cy3-AF647
MERGE
TGN46-HA
vs
TGN46-HA
TGN46-HA
vs
TGN46-GFP
TGN46-HA
vs
ST-GFP
TGN46-HA
vs
TGN46-shortTMD-GFP
TGN46-HA
vs
TGN46-ST TMD-GFP
B
Spearman coefficient A
Spearman coefficient B
0.8
0.6
0.4
0.2
0.0
-0.2
-0.4
ns
*
**
TGN46-HA TGN46-HA
TGN46-HA TGN46-GFP
TGN46-HA ST-GFP
TGN46-HA TGN46-shortTMD-GFP
TGN46-HA TGN46-ST TMD-GFP

**Figure S4. STORM analysis of intra-Golgi localization of different TGN46 mutants. (A)** Representative dual-color STORM images of TGN46-HA, labelled using AlexaFluor 405-AlexaFluor 647 (AF405-AF647) conjugated secondary antibodies (green channel), together with TGN46-HA, TGN46-GFP, ST-GFP, TGN46-shortTMD-GFP, or TGN46-ST TMD-GFP, labelled using Cy3-AlexaFluor 647 (Cy3-AF647) conjugated secondary antibodies (magenta channel). Images were rendered using Insight3. Scale bars are 2 µm. **(B)** Plots of the Spearman coefficient A (between green:magenta channels) and B (between magenta:green channels). Distributions contain information from > 5 cells from each of n=3 independent experiments; individual values are shown, with mean ± stdev.

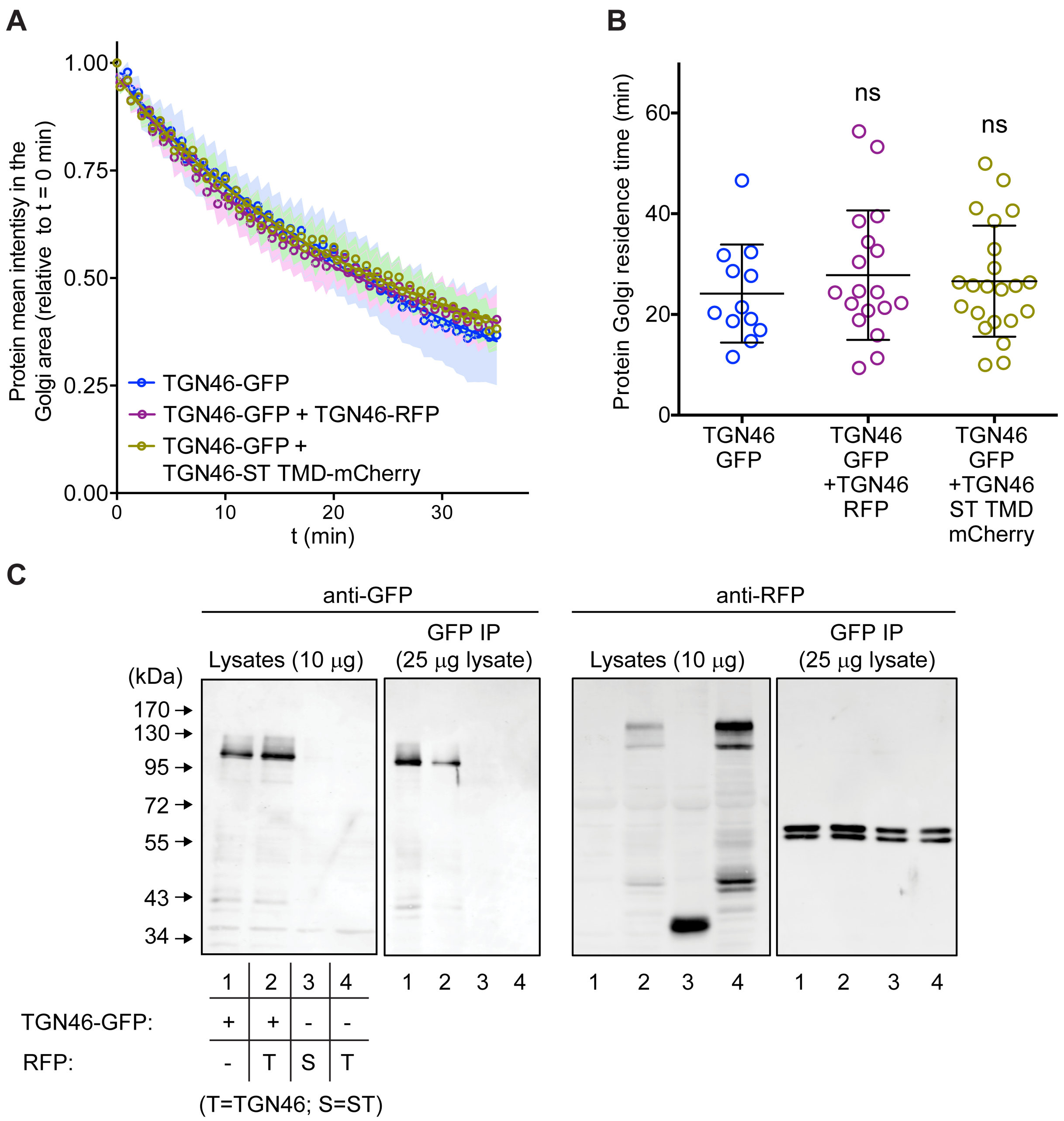


**Figure S5. Co-expression of different TGN46 proteins does not affect CARTS biogenesis or cargo export rate. (A)** Relative fluorescence intensity average time trace (mean ± s.e.m.) of FLIP experiments for TGN46-GFP obtained in cells expressing the different indicated proteins. Symbols correspond to actual measurements, solid lines to the fitted exponential decays. **(B)** Residence time in the perinuclear area measured as the half time of the FLIP curves shown in **(A)**. Results are from from 4–10 cells from each of n=3 independent experiments (individual values shown, with mean ± stdev). **(C)** TGN46-GFP and TGN46-mRFP do not co-immunoprecipitate. HeLa cells expressing TGN46-GFP (lanes 1), TGN46-GFP and TGN46-mRFP (lanes 2), ST-mCherry (lanes 3), or TGN46-mRFP (lanes 4), were lysed and part of the lysate was immunoprecipitated with an anti-GFP antibody. Both the lysate and immunoprecipitated fraction were analyzed by SDS-PAGE and Western blotting with anti-GFP and anti-RFP antibodies.

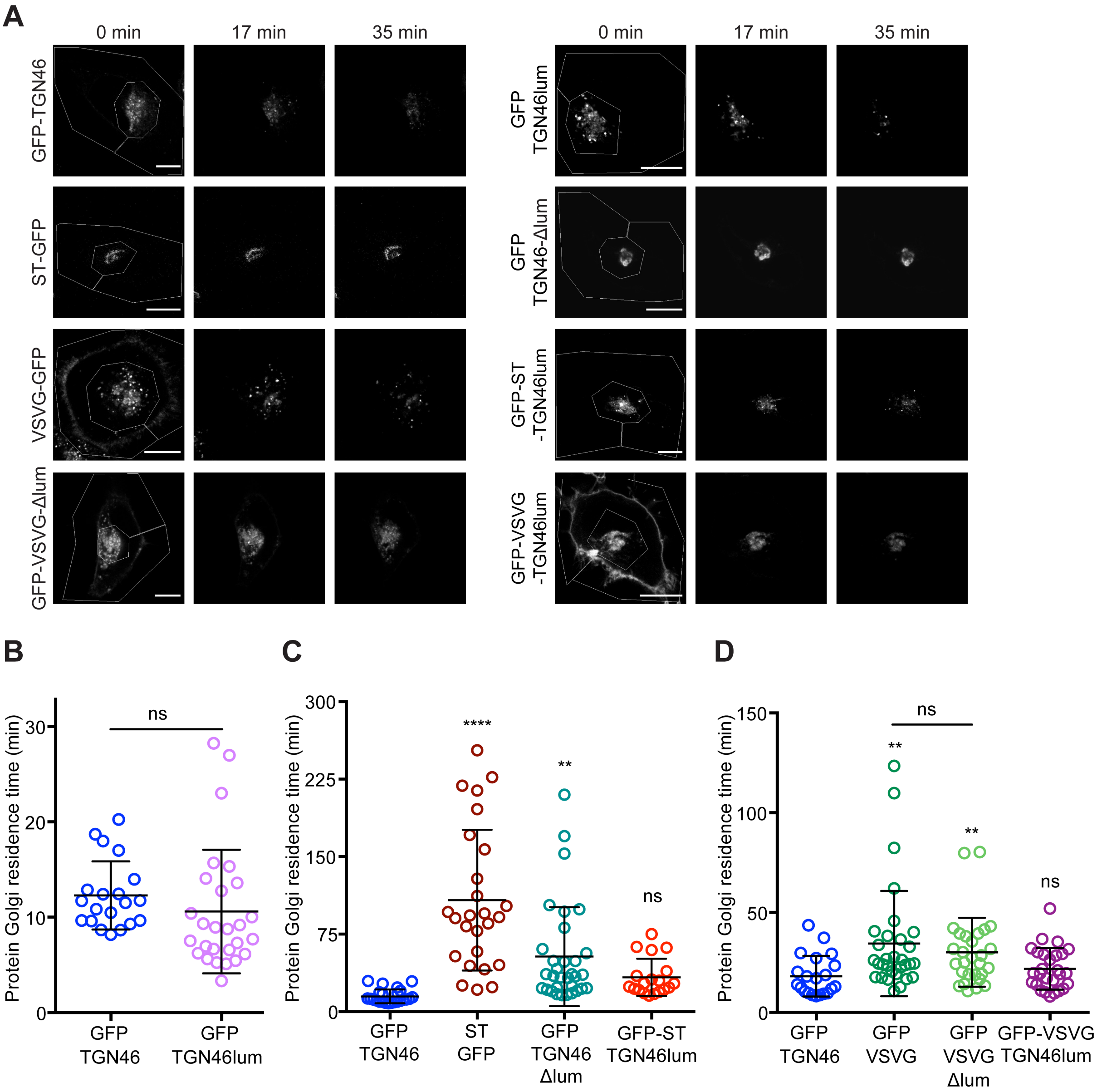


**Figure S6. The lumenal domain of TGN46 is necessary and sufficient for its CARTS-mediated export from the TGN. (A)** Fluorescence microscopy images obtained from a characteristic FLIP experiment assessing the export rate of the denoted proteins from the perinuclear area in HeLa cells. Time from the beginning of the FLIP experiments is indicated, and the area enclosed by the white lines shown in the left images denotes the photobleached area. Scale bares are 10 µm. **(B–D)** Residence time in the perinuclear area measured as the half time of the FLIP curves. Results are from 7–12 cells from n=3 independent experiments (individual values shown, with mean ± stdev). In **(B)**, unpaired two-tailed t test (ns, $p > 0.05$).

A

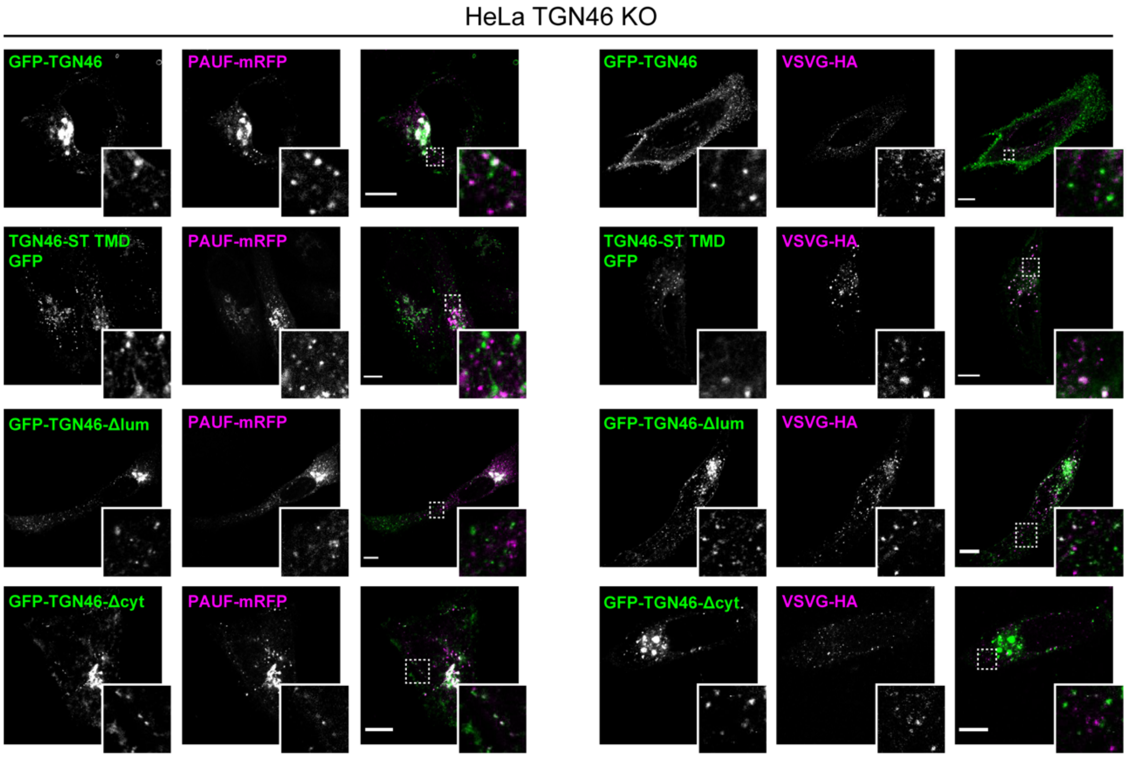


**Figure S7. The cargo sorting function of TGN46 is mediated by its lumenal domain. (A)** HeLa TGN46-KO cells co-expressing the different indicated proteins (green and magenta channels) were fixed, processed for immunostaining when required, and the localization of those proteins was monitored by fluorescence confocal microscopy. Insets correspond to zoom-in areas of the dashed, white boxed areas. Scale bars are 10 µm.

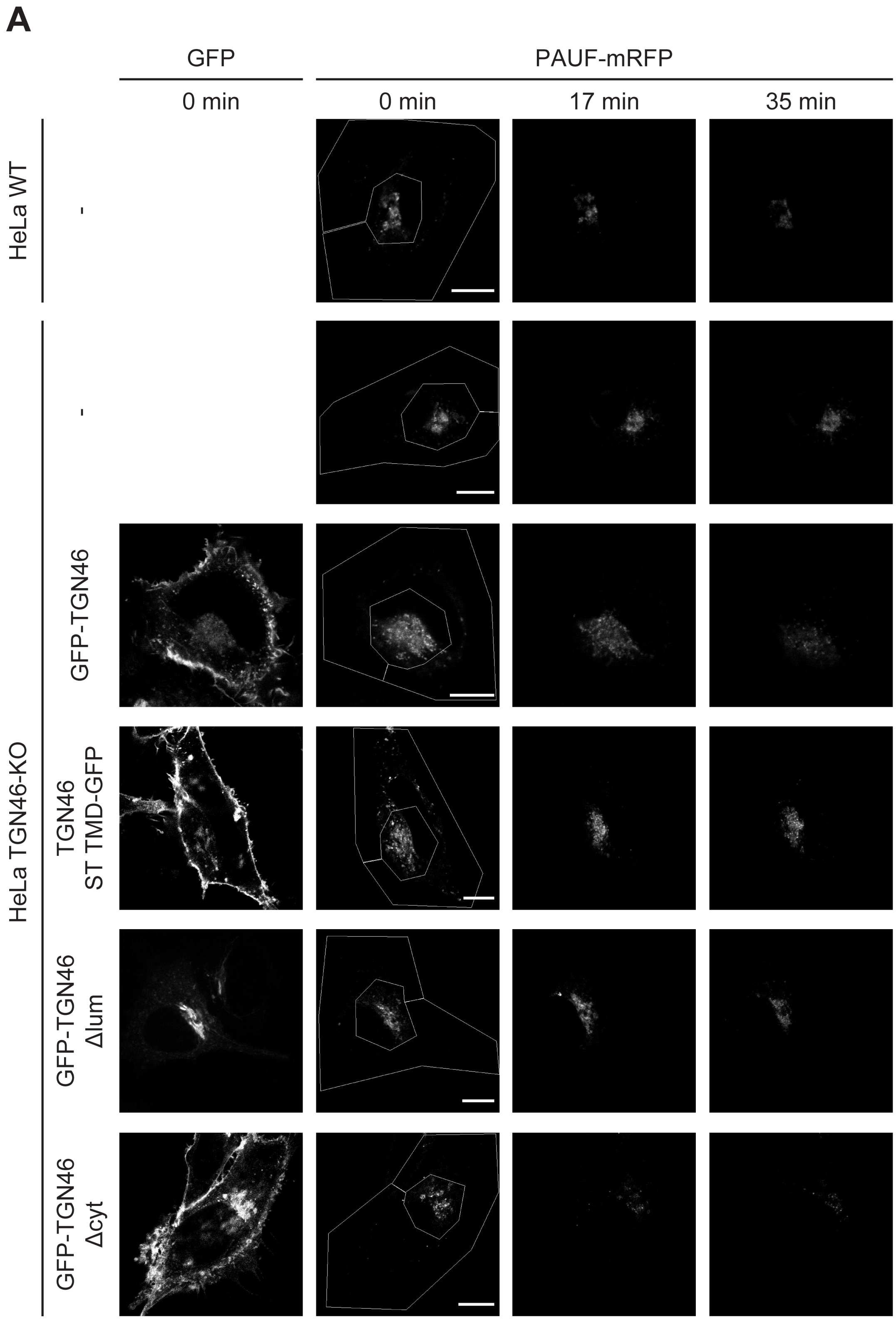


**Figure S8. Golgi export of PAUF-mRFP is dependent on the lumenal domain of TGN46. (A)** Fluorescence microscopy images obtained from a characteristic FLIP experiment assessing the export rate of PAUF-mRFP from the perinuclear area in HeLa cells (either WT or KO) expressing the indicated proteins. Time from the beginning of the FLIP experiments is indicated, and the area enclosed by the white lines shown in the left images denotes the photobleached area. Scale bars are 10 µm.

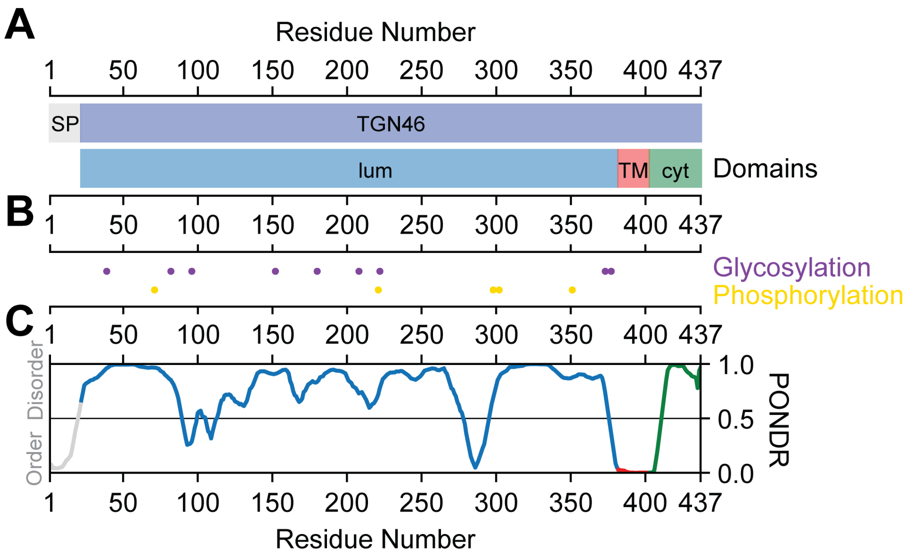


**Figure S9. TGN46 domain structure and sequence characteristics. (A)** On the top, signaling peptide region (SP, grey) and mature TGN46 (TGN46, blue). On the bottom, the lumenal (lum, blue), transmembrane (TM, red) and cytoplasmic (cyt, green) domains. **(B)** Post-translational modifications. On the top, the glycosylation sites (purple) and on the bottom, the phosphorylation sites (yellow). **(C)** Disorder propensity indicated by PONDR score using the VL-XT algorithm (Romero et al., 1997; Li et al., 1999). Values ranging 0.0-0.5 indicate order and values ranging 0.5-1.0 indicate disorder.
